# Retinal neuroblast migration and laminar organization requires the cytoskeletal-interacting protein Mllt11

**DOI:** 10.1101/2022.03.01.482481

**Authors:** Marley Blommers, Danielle Stanton-Turcotte, Angelo Iulianella

## Abstract

**Background:** The vertebrate retina is an organized laminar structure comprised of distinct cell types populating three nuclear layers. During development, each retinal cell type follows a stereotypical temporal order of genesis, differentiation, and migration, giving rise to its stratified organization. Once born, the precise positioning of cells along the apico-basal (radial) axis of the retina is critical for subsequent connections to form, relying on orchestrated migratory processes. While these processes are critical for visual function to arise, the regulators of cellular migration and retinal lamination remain largely unexplored.

**Results:** We report a role for a microtubule-interacting protein, Mllt11 (Myeloid/lymphoid or mixed-lineage leukemia; translocated to chromosome 11/All1 Fused Gene From Chromosome 1q) in mammalian retinal cell migration during retinogenesis. We show that *Mllt11* loss-of-function in mouse retinal neuroblasts affected the migration of ganglion and amacrine cells into the ganglion cell layer, and led to their ectopic accumulation in the inner plexiform layer.

**Conclusions:** We demonstrate that Mllt11 plays a critical role in the migration and lamination of neurons in the retina, and its loss impacted formation of the basal-most retinal layers.

## INTRODUCTION

The vertebrate retina is a specialized extension of the brain responsible for the detection, processing, and encoding of visual information. Its function arises from a laminated cellular architecture, comprised of seven major cell types organized into three layers of cell bodies, or nuclear layers, and separated by two synaptic, or plexiform layers ^1,2^. The apical, or outermost layer (outer nuclear layer; ONL), contains cell bodies of photoreceptors (rods and cones)^2^. The basal, or innermost layer (retinal ganglion cell layer; GCL) contains cell bodies of retinal ganglion cells (RGCs), which are the output cells of the retina, relaying visual information to the brain for processing via their collection of axons forming the optic nerve. Bipolar, amacrine, and horizontal cells all have cell bodies located in the inner nuclear layer (INL) between photoreceptor and ganglion cells. Bipolar cells form connections with photoreceptors in the outer plexiform layer (OPL), and with ganglion cells in the inner plexiform layer (IPL)^2^. Additional lateral processing exists in the OPL and IPL by horizontal and amacrine cells, respectively.

The arrangement of neuronal cell bodies into distinct layers with respect to their function is called retinal lamination^3,4^. Like the neocortex, disorganized lamination in the retina results in disrupted synaptic connectivity and impaired function^4^. For a layered structure to arise, retinal neurons are derived from a pool of multipotent retinal progenitor cells (RPCs) in a distinct, although overlapping, birthing order as they migrate basally to occupy their respective nuclear layer^5^. In the mouse, retinal progenitors are neatly arranged in a germinal zone known as the neuroblastic cell layer (NBCL)^6^. Within this layer, RPCs follow a stereotypic sequence of cell cycle exit, cell-type determination, migration, and differentiation to give rise to the cellular diversity that exists within the retina^5^. As early as embryonic day 10 (E10) in the mouse retina, neural progenitors from the germinal region begin to differentiate to the earliest born cell type, the ganglion cells, and migrate basally to form the putative GCL, separated from the underlying NBCL by a narrow gap; the putative IPL where contacts will form between ganglion, bipolar, and amacrine cells^4,6,7^. At E11 some neural stem cells in the NBCL begin to differentiate into horizontal and amacrine cells, the next born cell types^6^. In the mouse, retinal neurogenesis persists postnatally, with rod photoreceptor and bipolar cell genesis peaking at birth and during the first postnatal week, respectively^8^. Once differentiated, the proper positioning of postmitotic retinal neurons along the apico-basal (radial) axis of the retina is crucial for constructing the laminar architecture, and for successive connections to form. The formation of functional neuronal circuitry relies on precise positioning of cells in their respective nuclear layers; thus, the migration, lamination, and positioning of cells along the apico-basal axis sets the groundwork for subsequent visual function to arise. Despite its functional importance, the migratory mechanisms underlying retinal neurogenesis have not been fully explored.

Retinal ganglion cells are the first cells to differentiate and migrate away from their apical birthplace to occupy the most basal retinal layer adjacent to the lens, from E10 to postnatal day 0 (P0), with peak neurogenesis levels at E14.5^6,9^. RGCs migrate away from the apical surface via a bipolar somal translocation mechanism in which the soma moves basally while remaining anchored to both apical and basal surfaces through cytoskeletal elements^9^. In instances where microtubule or cytoskeletal integrity is impaired, RGCs may unfavorably adopt a multipolar migration mode, which is less efficient and result in ectopically placed cells^10^. Since the retina relies on serial connections to form the visual pathway, displacement of cells from their respective layer during early development likely disrupts circuitry formation and function.

In an effort to identify regulators of neural migration, we previously reported the expression of Mllt11/Af1q (Myeloid/lymphoid or mixed-lineage leukemia; translocated to chromosome 11/All1 Fused Gene from Chromosome 1q) in the developing central nervous system, including robust expression in the retina^11^. Mllt11 is a vertebrate-specific 90 amino acid protein first identified in an acute myelomonocytic leukemia carrying the t(1;11)(q21;q23) translocation that creates an abnormal protein fused to the chromatin remodelling factor Mll^12^. Recent work from our lab has shown that Mllt11 is a microtubule-associating protein that is required for cortical neuron migration and neuritogenesis^13^. However, its role in retinal histogenesis is unknown. Using a conditional inactivation approach, we reveal a role for Mllt11 in the formation of a laminated retinal neuroanatomy by regulating neuroblast migration into the GCL.

## RESULTS AND DISCUSSION

### Conditional Knockout of *Mllt11* in Migrating Neuroblasts During Retinogenesis

Mllt11 is expressed in developing neurons of the central nervous system, including those that comprise the retina, but its role in retinogenesis is unknown. Yamada et al. (2014)^11^ first described its expression pattern in the vertebrate eye, revealing it to be spatially restricted to cells comprising the retinal basal layers (INL and GCL) and their axons, which collectively make up the optic nerve. At the developmental time points reported in that study (E13.5 and E16.5), retinal cells are rapidly proliferating, differentiating into ganglion, amacrine, and horizontal cells, and migrating away from the RPC pool to their basal destination. Due to its spatial and temporal specificity, *Mllt11* may play a role in the migration of post-mitotic neurons and lamination of the retina.

To investigate this, we utilized a loss-of-function approach by generating mice carrying a null mutation in the *Mllt11* gene. This approach would allow for the proper characterization of its role in the developing neural retina. The *Cux2iresCre* transgenic line served as an appropriate model to delete *Mllt11* in Cux2-expressing cells during retinogenesis, while simultaneously activating fluorescent reporter transgene *tdTomato*, allowing for optimal visualization of mutant cells. Previous work from our lab demonstrated that Cux2 regulates neurogenesis^14–17^, and Cux2 is expressed in the neural retina during development, including migrating neuroblasts (Figure 1, see below). Our genetic ablation strategy involves using *Cux2iresCre* to inactivate a floxed *Mllt11* allele to produce cKO mice, which lack *Mllt11* expression in migratory neuroblasts and basal layers of the retina. At the same time, we used the Ai34 reporter allele, which expresses *tdTomato* following Cre recombinase activity to reveal *Cux2*-positive cells in the developing retina^13^. We used this transgenic model to effectively characterize the role of *Mllt11* in the migration, lamination, and histogenesis of the neural retina.

**Figure 1:**
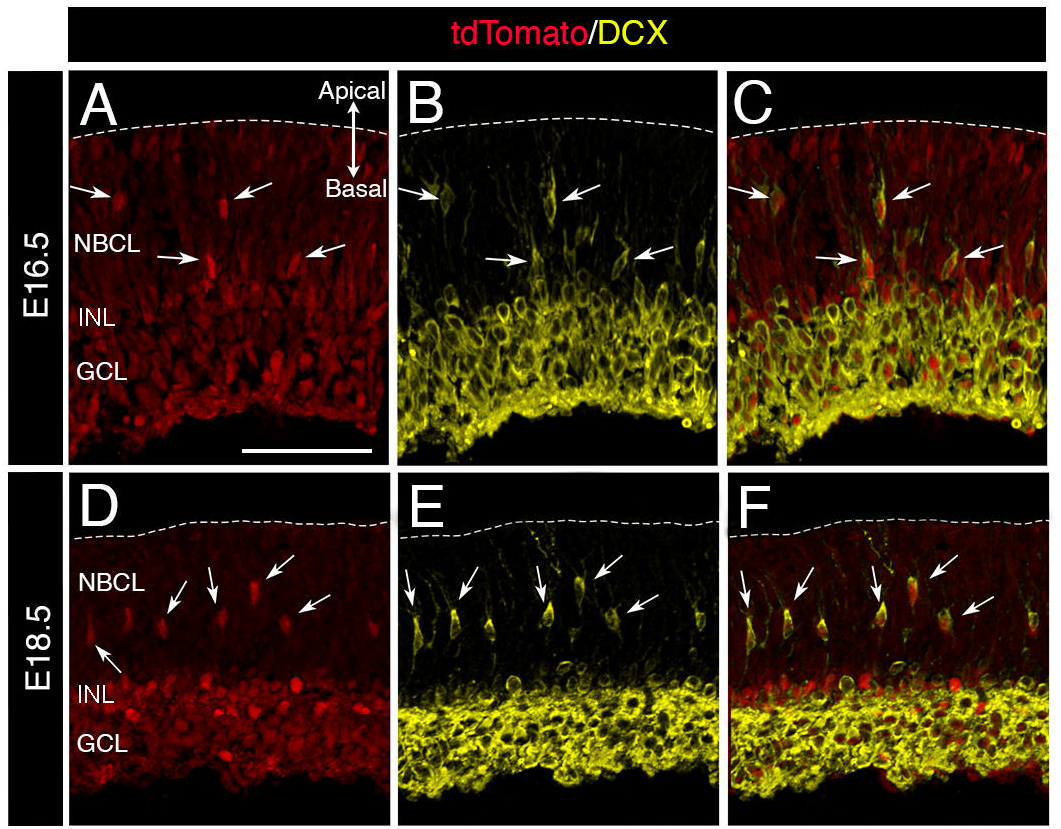
TdTomato and DCX co-expression revealed *Cux2iresCre*-mediated deletion in migrating retinal neuroblasts. (A-F) Sagittal slices of the retina at E16.5 and E18.5 stained for migratory neuroblast marker DCX and fluorescent reporter TdTomato. Arrows indicate co-expression of DCX and TdTomato in migratory neuroblasts within the NBCL. (A) E16.5 TdTomato fluorescence within the putative GCL and INL/IPL retinal layers and in migratory neuroblasts moving basally. (B) E16.5 DCX labelling of migratory neuroblasts in the NBCL (arrows) and putative GCL and INL (C) E16.5 Co-expression of TdTomato and DCX in migratory neuroblasts within the NBCL, and putative GCL and INL (arrows). (D) E18.5 TdTomato fluorescence within migratory neuroblasts in the NCBL (arrows), GCL and INL. (E) E18.5 DCX labelling of migratory neuroblasts in the NBCL, GCL and INL. (F) E18.5 Co-expression of TdTomato fluorescence and DCX in the same population of migratory neuroblasts in the NBCL (arrows). L. Scale bar: 50μm for (A-F). GCL, ganglion cell layer; INL, inner nuclear layer; NBCL, neuroblastic cell layer.

We first confirmed *Cux2* activity in migrating retinal neuroblasts, revealed by the restricted TdTomato fluorescence expression in migrating neuroblasts within the cohort of Doublecortin (DCX)-positive cells in the NBCL and basal retinal layers (putative INL/IPL and GCL) at E16.5 (Figure 1A-C) and E18.5 (Figure 1D-F). The *Cux2* fate-mapped cells closely reflect the previously reported *Mllt11* mRNA expression pattern in the retina, signifying that the use of the *Cux2iresCre* driver is an appropriate approach to inactivate *Mllt11* in the migrating neuroblasts and basal layers of the neural retina.

### Loss of *Mllt11* in Migrating Neuroblasts Reduces Cellular Composition of the GCL

With an appropriate genetic model to conditionally delete *Mllt11* in migratory retinal neuroblasts, we set out to investigate its role in the process of retinogenesis. We used DAPI (4’,6-diamidino-2-phenylindole) nuclear staining at E16.5 and E18.5 to evaluate any gross phenotypic consequences due to the conditional-knockout (cKO) of *Mllt11* (Figure 2). At these time points, ganglion cell genesis, differentiation, and migration are well underway, along with moderate levels of horizontal, amacrine, and cone photoreceptor genesis (PMID: 29375289). *Mllt11* cKO mutants displayed fewer DAPI+ nuclei within the GCL compared to wild-type littermate controls at both E16.5 (Figure 2A-C) and E18.5 (Figure 2D-F). The GCL could be readily identified by the narrow gap (putative IPL) separating it from the underlying NBCL. The NBCL in both mutants and controls were densely packed with progenitors and appeared comparable between controls and cKOs. However, DAPI+ nuclei within the GCL were significant reduced in *Mllt11* mutants at both E16.5 (Figure 2C; WT = 33.92 +/- 1.34, cKO = 29.19 +/- 0.92, P = 0.002, N = 4, Welch’s t-test) and E18.5 (Figure 2F; WT = 29.10 +/- 2.59, cKO = 23.72 +/- 0.79, P = 0.008, N = 5, Welch’s t-test), with tissue gaps clearly visible in the mutant GCL.

**Figure 2:**
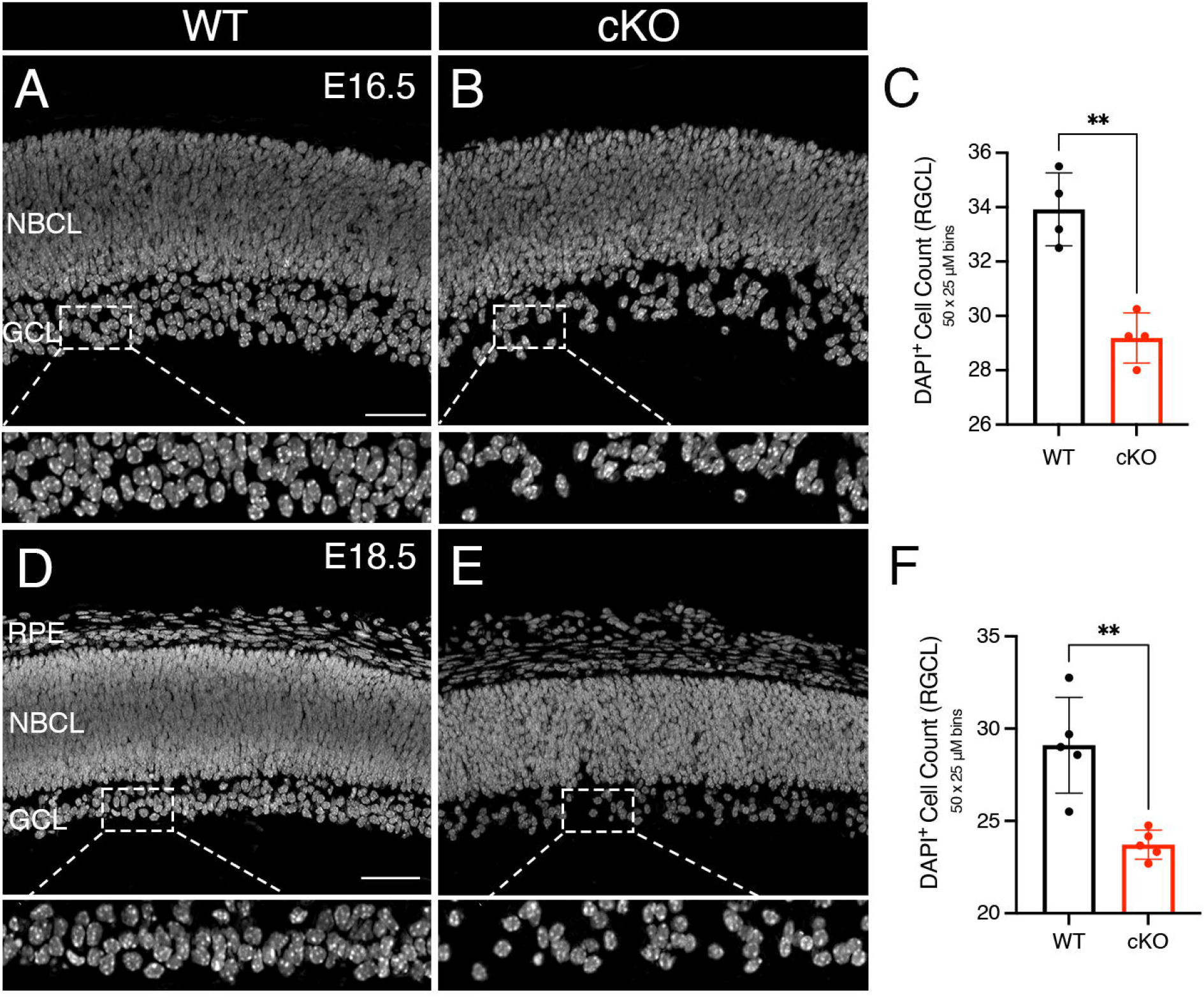
*Mllt11* loss resulted in fewer cells populating the GCL. (A, B, D, E) Sagittal sections of the retina at E16.5 and E18.5 stained with the nuclear marker DAPI. Corresponding images beneath each panel show cells occupying the GCL zoomed in. (A, B) E16.5 WT controls (A) had significantly more cells within the GCL compared to E16.5 cKO (B). (C) Bar chart comparison of WT controls vs. Mllt11 cKO mutant total cell counts within the GCL, 50μm x 25 μm bins for (A and B). (D-E) E18.5 WTs (D) had significantly more cells within the GCL compared to E18.5 cKOs (E). (F) Bar chart comparison of WT vs. cKO total cell counts within the GCL, 50μm x 25 μm bins for (D and E). Welch’s t-test, (A, B) N =4, (D, E) N = 5. Data presented as mean ± SD. n.s., not significant. **P≤0.01. Scale bar: 50μm for (A, B, D, E). GCL, ganglion cell layer; NBCL, neuroblastic cell layer; RPE, retinal pigmented epithelium.

### Loss of *Mllt11* Leads to Impaired Lamination and Positioning of Cells in the GCL

To confirm the identity of the cells in the presumptive GCL, we used immunohistochemistry against RNA Binding Protein with Multiple Splicing (RBPMS) to robustly label ganglion cells in the GCL of the vertebrate retina^18^. In support of our DAPI findings, RBPMS staining revealed reduced cellular density in the GCL of *Mllt11* cKO mutants compared to WT controls at E16.5 (Figure 3A-E) and E18.5 (Figure 3G-K). High magnification views of the RBPMS staining revealed reduced cellular density in the GCL of *Mllt11* mutant at E16.5 (Figure 3C) and E18.5 (Figure 3I). Densitometric analysis of RBPMS staining revealed more dispersed cellular packing and organization within the GCL of cKOs at E16.5 (Figure 3F; WT average density = 206.4 +/- 12.11, N = 4, cKO average density = 185.5 +/- 11.81, N = 5, P = 0.037, Welch’s t-test) and E18.5 (Figure 3L; WT average density = 244.9 +/- 15.48, cKO average density = 187.0 +/- 11.44, P = 0.001, N = 4, Welch’s t-test).

**Figure 3:**
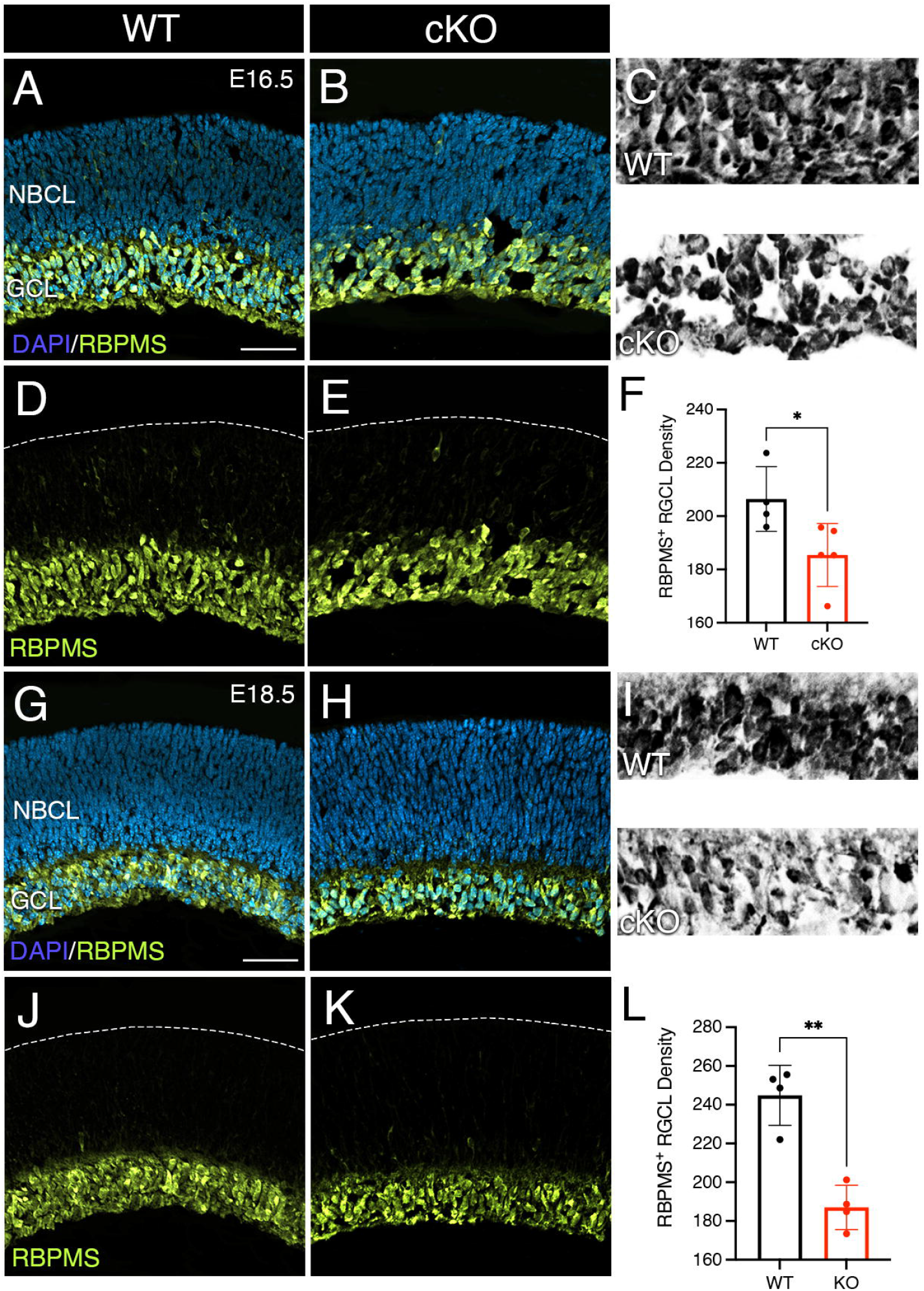
*Mllt11* loss leads to reduced RBPMS+ cell counts in the GCL. (A-E) Sagittal sections of the WT (A, D) and cKO (B, E) retina at E16.5 stained with RBPMS, labeling RGCs, counterstained with DAPI (A, B). (C) Zoomed in images of RBPMS staining in the GCL of E16.5 WT (top panel) and cKO (bottom panel), converted to a binary image to highlight reduced cellular density in *Mllt11* mutant GCLs. (F) Bar graph of density measurements taken of the GCL in 50μm x 25 μm bins revealed significant decreases density in cKOs compared to WTs at E16.5. (G-K) RBPMS staining in E18.5 retinas of WT (G, J) and *Mllt11* cKO mutant (H, K), counterstained with DAPI (G, H). (I) Zoomedin images of RBPMS staining in the GCL of E18.5 WT (top panel) and cKO (bottom panel), converted to a binary image to highlight reduced cellular density in *Mllt11* mutant GCLs. (L) Bar graph of density measurements taken of the GCL in 50μm x 25 μm bins revealed significant decreases density in cKOs compared to WTs at E18.5. Welch’s t-test, (A, B, D, E) N =4, (G, J) N = 4, (H, K) N =5. Data presented as mean ± SD. n.s., not significant. *P≤0.05; **P≤0.01; ***P≤0.001; ****P≤0.0001. Scale bar: 50μm for (A, B, D, E, F, H, J, K). GCL, ganglion cell layer; NBCL, neuroblastic cell layer.

The GCL consists of a heterogeneous group of neurons, including both ganglion cells and displaced amacrine cells^19^. While most amacrine cells populate the INL, there are amacrine cells that are displaced in basal boundary of the GCL, which constitute a large percentage of neurons in this layer. In mice displaced amacrine cells constitute about 55-56% of the neurons found in the GCL^20^. While RBPMS served as a useful marker for RGCs of the GCL, it does not capture displaced amacrine cells, making any conclusions regarding the cellular composition of the GCL challenging without an alternative marker.

Sox2 is a marker of both neural progenitors constituting the NBCL, distinguishing it from the RBPMS-labelled GCL, as well as a subset of amacrine cells^21^. At E18.5 Sox2 was expressed throughout the neuroblastic layer as well as in a subset of cells occupying the INL and the GCL in the embryonic retina (Figure 4A, C). Lin et al. (2009)^21^ demonstrated that the ectopic expression of Sox2 in cells outside the NBCL induced amacrine cell differentiation and cell cycle exit. Sox2 immunostaining can therefore be used to detect displaced amacrine cells within basal surface of the GCL (arrows, Figure 4C). At E18.5, *Mllt11* cKOs had fewer Sox2+ cells within the GCL and more within the INL/IPL compared to controls (Figure 4A,B; arrows in Figure 4C, D). The overall (INL/IPL + GCL) Sox2+ cell counts normalized over DAPI did not differ between genotype, however, their distribution across retinal layers did. Specifically, cKOs had more Sox2+ cells in the INL and IPL (position 1, Figure 4E, Figure 4F; WT = 17.83 +/- 1.9, cKO = 26.90 +/- 3.84, P = 0.036, N =3, Welch’s t-test) and significantly fewer in the basal-most edge of the GCL (position 2, Figure 4E, arrows in Figure 4D, Figure 4F; WT = 39.23 +/- 5.22, cKO = 15.77 +/- 2.05, P = 0.009, N = 3, Welch’s t-test). The shift in Sox2+ cell numbers within the INL/IPL of *Mllt11* mutants is likely due to the inability of these cells to properly migrate into the basal surface of the GCL and differentiate into displaced amacrine cells (Figure 4E, F). Overall, these findings suggest that Mllt11 may regulate the migration of amacrine and ganglion cells out of the IPL/INL region of the retina to settle into a displaced basal destination.

**Figure 4:**
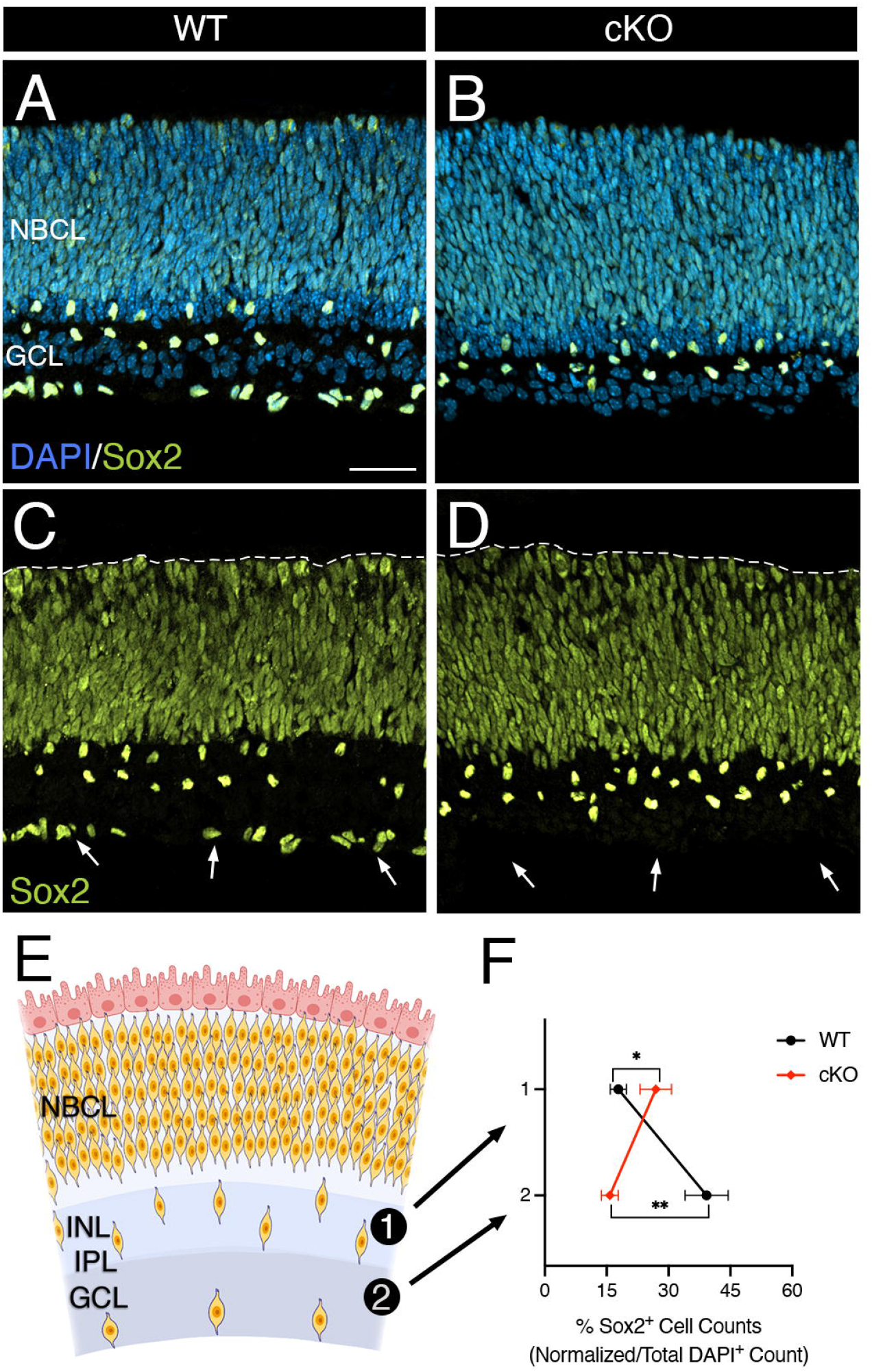
Sox2+ staining revealed impaired GCL formation in *Mllt11* cKO mutant retinas. (A-D) Sox2 staining in E18.5 sagittal retinal sections labeled the progenitor region (NBCL), presumed amacrine cells in the INL/IPL, and displaced amacrine cells in the GCL. (A-B) Sox2/DAPI co-staining revealed a band of Sox2+ cells at the basal edge of the GCL in WTs (A), which was absent in cKOs (B). (C-D) Sox2 staining in the displaced amacrine cells in the E18.5 WT GCL (arrows, C) was absent in the cKO retinas (arrows, D). (E) Schematic depiction of Sox2+ NBCL and the two regions of interest for comparing cellular composition: the INL/IPL region basal to the NBCL (area 1), and the basal edge of the GCL (area 2). (F) Line graphs of the relative distribution of Sox2+ cells in 125μm x 35μm bins sampled in areas 1 and 2. WTs had significantly more Sox2+ cells in the GCL (Area 2), whereas *Mllt11* cKOs had more ectopically placed cells caught in the INL/IPL region (Area 1). Welch’s t-test, (A-D) N =3. Data presented as mean ± SD. n.s., not significant. *P≤0.05; **P≤0.01; ***P≤0.001; ****P≤0.0001. Scale bar: 50μm for (A-D). GCL, ganglion cell layer; INL, inner nuclear layer; IPL, inner plexiform layer; NBCL, neuroblastic cell layer.

### *Mllt11* is Required for the Migration of Retinal Neuroblasts

To further investigate a migratory defect in *Mllt11* mutants, we used DCX to visualize migratory neuroblasts within the NBCL and cells within the GCL^22,23^. DCX is a cytosolic protein widely expressed in immature cells of the vertebrate retina and their processes as they migrate through the progenitor layer. DCX also labels a subset of cells occupying the GCL, including a subtype of RGCs and a population of horizontal cells that initially migrate all the way into the GCL before reverting their migratory course back to the INL^23^ (Figures 1 and 5). Relative to the entire thickness of the retina, the distance travelled by migratory DCX+ neuroblasts within the *Mllt11* cKO mutant NBCL was shorter and restricted to the apical retinal surface (Figure 5B, D; see below), compared to the more widespread distribution of DCX+ cells in WT controls at E18.5 (Figure 5A, C). *Mllt11* mutant neuroblast processes were shorter and did not have the morphology typical of wild type neuroblasts, with processes spanning the NBCL (Figure 5E, F; see below). This provides additional evidence in support of a migratory defect in *Mllt11* mutant retinas.

**Figure 5:**
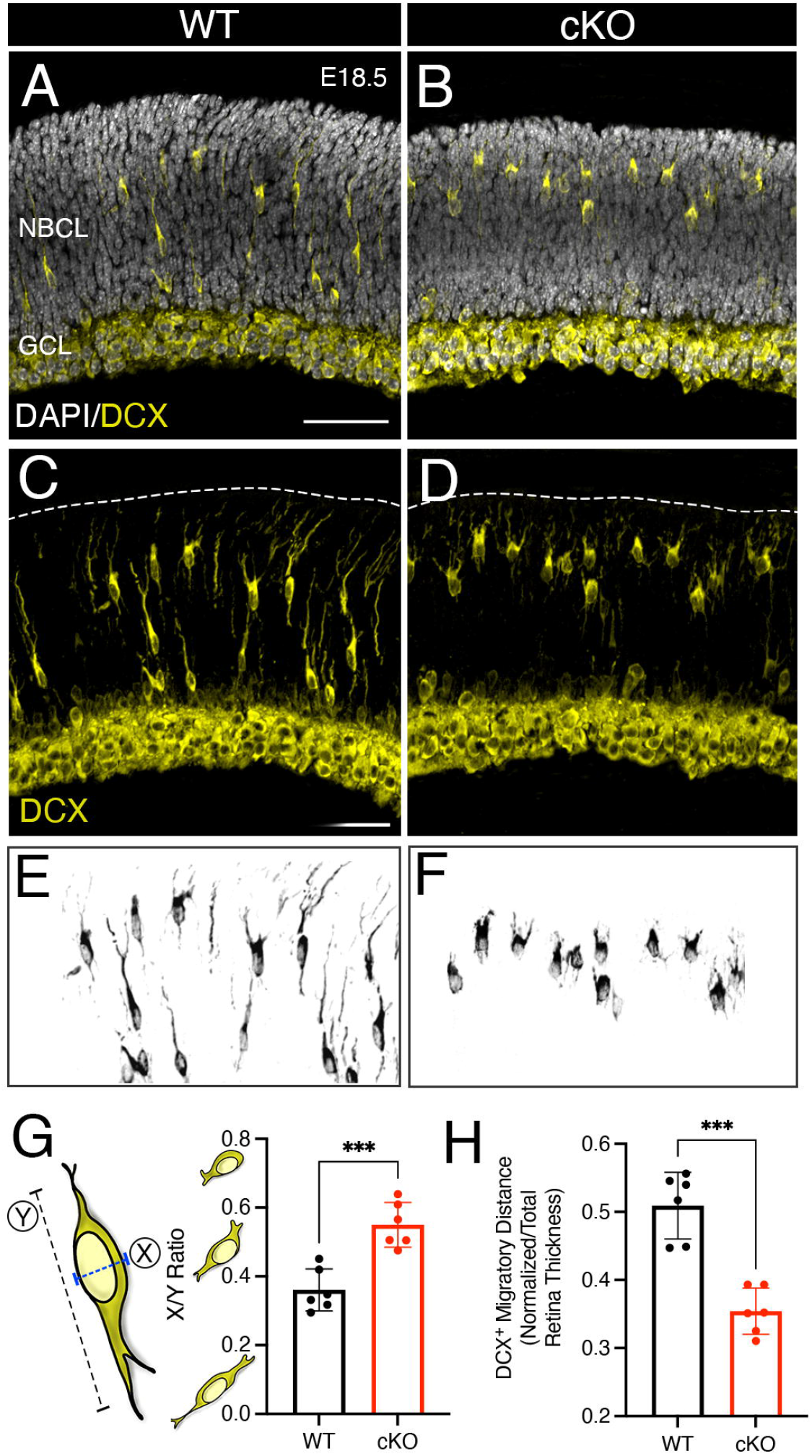
*Mllt11* loss resulted in reduced bipolar neuroblast morphology and impaired migration in the NBCL. (A-F) DCX staining of sagittal sections of the E18.5 retina labeled migratory neuroblasts within the NBCL and cells in the GCL, co-stained with DAPI (A, B). DCX+ migratory neuroblasts were aberrantly localized in the apical region of the NBCL of cKO retinas (D), compared to the more extensive distribution of neuroblasts throughout the NBCL of WTs (C). DCX staining reveals extensive bipolar cellular processes in migratory neuroblasts in the WT NBCL (C), *Mllt11* cKO mutant neuroblasts (D) displayed more rounded multipolar morphologies. (E-F) Converted binary image of DCX staining in the NBCL of a WT (E) and cKO (F) retina highlighting the differences in cellular morphology. (G) Schematic and bar graph of X/Y ratio measurements of DCX+ cells for morphometric analysis (X, width along the major axis; Y, length along major axis). WT migratory neuroblasts are more elongated as indicated by a lower X/Y ratio, while cKOs are more globular in shape with a ratio closer to 1. (H) Bar graph quantifying the average migratory distance of DCX+ neuroblasts, expressed as a ratio of the distance from the apical surface normalized to the entire thickness of the retina. WT neuroblasts were on average displaced further away from the apical surface vs cKOs. Welch’s t-test, (A-H) N =4. Data presented as mean ± SD. n.s., not significant. *P≡0.05; **P≡0.01; ***P≤0.001; ****P≤0.0001. Scale bar: 50μm for (A-D). GCL, ganglion cell layer; NBCL, neuroblastic cell layer.

To investigate a cellular mechanism underlying the *Mllt11* mutant migratory phenotype, we used a method of morphometric analysis previously described by Yu et al. (2013)^24^. We calculated an X/Y ratio (X being the width of the cell perpendicular to the major axis, and Y being the longest length of the cell along the major axis) for all DCX+ migratory neuroblasts contained within the NBCL, and then averaged them for each individual (Figure 5G). *Mllt11* mutant mice had an X/Y ratio closer to 1, indicative of round or globular cell morphology with minimal projections extending out from the cell body, whereas WT mice had an X/Y ratio closer to 0, indicative of a long, filamentous cellular morphology with extensive processes emanating from the cell body (Figure 5G, WT X/Y ratio = 0.36 +/- 0.06, cKO X/Y ratio = 0.55 +/- 0.06, P = 0.0004, N = 6, Welch’s t-test). These results confirm the difference in neuroblast morphologies in *Mllt11* mutants vs. controls, with the former displaying a less elongated bipolar organization and instead displayed shortened multipolar processes (Figure 5C-F).

Migrating ganglion cells typically adopt a bipolar somal translocation mechanism when migrating through the NBCL toward the basal GCL for proper positioning and lamination^9^. This type of elongated bipolar morphology is clearly outlined by DCX-staining in the NBCL in the control WT retinas (Figure 5C, E). However, the loss of *Mllt11* led to neuroblasts adopting a multipolar morphology (Figure 5D, F), which likely contributed to the aberrant positioning of DCX+ cells more apically in the developing retina due to reduced migratory ability of the cells (Figure 5H, WT average normalized migration distance = 0.51 +/- 0.05, cKO average normalized migration distance = 0.35 +/- 0.03, P = 0.0001, N = 6, Welch’s t-test). These findings suggest that *Mllt11* mutants have displaced or ectopic retinal neurons principally due to a disruption in their cellular projections and attachment points to the apical and INL boundary regions. We recently showed that Mllt11 interacts with the microtubule cytoskeleton in developing cortical neurons ^13^. This is consistent with a retinal neuroblast migration phenotype in *Mllt11* cKO retinas, leading to reduced numbers of ganglion and amacrine cells occupying the GCL.

To confirm a migratory defect in Mllt11 mutant retinas, we performed an EdU (5-ethynyl-2’-deoxyuridine) birth dating experiment, pulsing dams at E14.5 to capture RGC peak neurogenesis, and analyzed retinas at E18.5. This allowed us to follow the origin and displacement of retinal neurons from their apical birthplace to their destination in the GCL, and evaluate whether *Mllt11* loss impacted the final laminar distribution of RGCs during development. An E14.5 pulse of EdU labeled cells within the GCL of E18.5 WT retinas (Figure 6A, arrows in Figure 6C). *Mllt11* cKOs displayed a clear reduction of EdU+ cells populating the GCL, and instead showed ectopic EdU+ cells arrested within the IPL (Figure 6B, arrows, Figure 6D). The distribution of EdU+ cells differed significantly between WT and cKO mice, with WT mice having a greater proportion of EdU+/DAPI+ cells in the GCL compared to mutants (WT = 29.1 +/- 2.8, cKO = 16.8 +/- 8.9, P = 0.0005, N = 5, Welch’s t-test; Figure 6F), whereas mutants had a greater proportion of EdU+/DAPI+ cells in the INL/IPL region (WT = 19.1 +/- 2.8, cKO = 31.1 +/- 6.7, P = 0.006, N = 5; Figure 6F). The EdU birth dating experiment revealed that *Mllt11* loss impacted the distribution of cells within the basal layers of the developing retina. The results confirmed the severe reduction of displaced Sox2+ amacrine cells in the basal edge of the mutant GCL (Figure 4), likely due to altered neuroblast morphologies indicative of reduced migratory potential (Figure 5).

**Figure 6:**
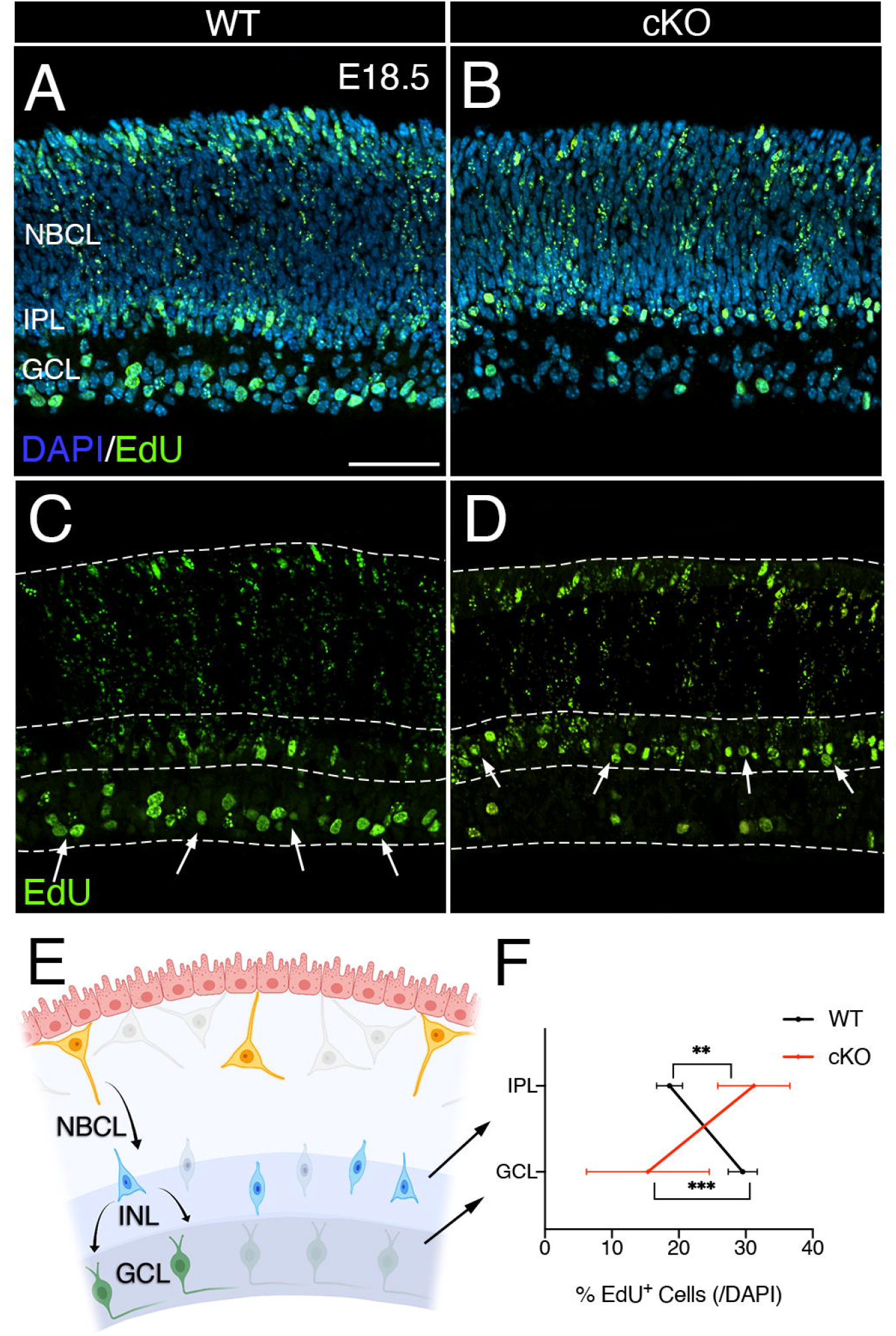
EdU birth dating demonstrated reduced migration of RGCs in the *Mllt11* cKO GCL. (A-D) Sagittal sections of E18.5 retinas following an EdU pulse at E14.5 to label RGCs. *Mllt11* cKOs, co-stained with DAPI (A, B). (C, D) EdU+ nuclei labeling revealed a larger number of EdU+ cells occupying the WT GCL (arrows, C), compared to cKOs, which instead were ectopically localized to the IPL (arrows, D). (E) Schematic representation of neuroblast (yellow) migration from the apical surface (RPE, pink), passing through the putative INL (blue), ultimately populating the GCL (green). (F) Line graph representing the relative distribution of EdU+ cells normalized over total DAPI cells in WT vs. *Mllt11* cKO in 150μm x 25μm (INL/IPL) and 150μm x 25μm (GCL) bins. WTs had more EdU+ cells within the GCL, whereas cKOs had more in the IPL, suggesting a migratory defect. Welch’s t-test, (A-H) N =4. Data presented as mean ± SD. n.s., not significant. *P≤0.05; **P≤0.01; ***P≤0.001; ****P≤0.0001. Scale bar: 50μm for (A-D). GCL, ganglion cell layer; INL, inner nuclear layer; IPL, inner plexiform layer; NBCL, neuroblastic cell layer; RPE, retinal pigmented epithelium.

Visual function relies on the proper migration and lamination of retinal neurons into their respective layers, which is necessary for connections to form between them. Crucial to our understanding of retinal histogenesis requires a genetic and cellular analysis of the regulatory events controlling neuroblast migration. Icha et al. (2016)^10^ described a switch to a multipolar migration mode often results in misplaced RGCs, with later-born cell types often following suit with ectopic displacement within the retinal layers. Using cell-type markers, EdU labeling, and cell morphometrics, we now provide evidence of the role of Mllt11, which interacts with the neuronal cytoskeleton^13^, in RGC and amacrine cell migration and proper lamination of the GCL. *Mllt11* regulates bipolar neuroblast morphology typical of migratory retinal cells, and in its absence DCX+ retinal cells adopted a multipolar morphology and failed to contribute to a proper laminated GCL.

Our findings add to a short list of cytoskeletal regulators implicated in retinal histogenesis. For example, the mutation of microtubule-associated protein like 1 (*Em11*) resulted in ectopic placement and disrupted lamination of retinal neurons (bipolar cells) during development^25^. Avilés et al. (2022)^26^ recently reported a role for the Fat-related protein 3 (Fat-3), which binds to various cytoskeletal regulators, in amacrine cell migration and laminar positioning. Our work adds Mllt11 to the list of cytoskeletal-interacting proteins that regulate retinal neuroblast migration and retinal laminar organization. Our recent proteomic findings have revealed that Mllt11 interacts with tubulins and atypical (non-muscle) myosins^13^. While it currently is unclear whether Mllt11 regulates microtubule stability or turnover, the association with myosins adds an intriguing possibility that Mllt11 may be involved in cytoplasmic transport requited for neurite extension and/or signalling receptor localization at leading processes.

*Mllt11* loss in retinal rneuroblasts led to severe alterations in cellular morphology, with neurites no longer spanning the extent of the NBCL (Figure 5). Our working model (Figure 7) is that Mllt11 is required for cytoskeletal stabilization of the growth cone, and *Mllt11* mutant cells lack robust cytoplasmic extensions projecting to the apical and basal edges of NBCL, which prevent translocation from the apical surface to the NBCL-GCL boundary during retinal development. Future work will focus on the underlying regulatory interactions between Mllt11 and cytoskeletal proteins important for retinal neuroblast migration.

**Figure 7:**
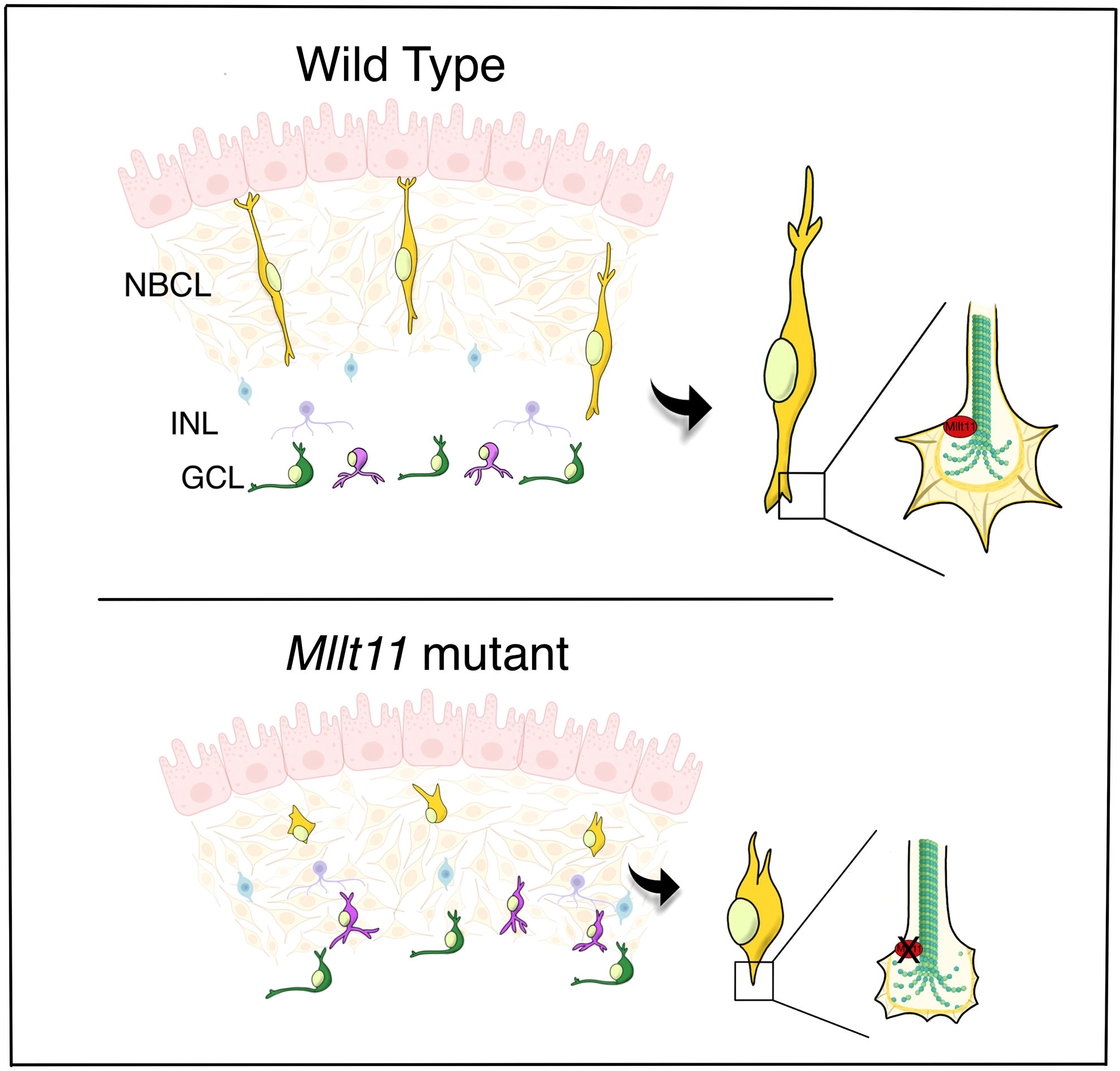
Model of Mllt11 function in migrating retinal neuroblasts. (Left) Wild type retinal neuroblasts (yellow) within the NBCL exhibit extensive cellular processes, which serve as apical and basal attachment points during migration. Proper positioning of RGCs (green) and displaced amacrine cells (purple) into the basal GCL require Mllt11 to interact with cytoskeletal elements. The interaction between Mllt11 and the cytoskeleton may regulate the ability of neuroblasts to adopt effective migratory mechanisms, such as the bipolar somal translocation, allowing for the formation of an organized and laminated retina. (Right) *Mllt11* mutant retinal neuroblasts within the NBCL (yellow) exhibit shortened and disrupted multipolar cellular processes. Adopting this aberrant morphology led to shortened translocation from the apical surface, resulting in RGCs (green) and displaced amacrine cells (purple) caught within apical layers (INL/IPL) in the *Mllt11* mutant retina. GCL, ganglion cell layer; INL, inner nuclear layer; NBCL, neuroblastic cell layer.

## EXPERIMENTAL PROCEDURES

### Animals

All animal experiments were done according to approved IACUC protocols at Dalhousie University. The *Cux2iresCre; Rosa26r^tdTomato/tdTomato^* wild type (WT) and *Cux2iresCre; Rosa26r^tdTomato/tdTomato^; Mllt11^flox/flox^* conditional knock-out (cKO) strains were generated to conditionally knock out Mllt11^13,17,27^. The Ai34 *Rosa26tdTomato* reporter mouse allowed for visualization of cells that had undergone Cre-mediated excision of a *floxed* translational stop sequence engineered upstream of the *tdTomato* cDNA within the ubiquitously expressed *Rosa26* locus.

### Histology

Eyes were dissected out and fixed in 4% paraformaldehyde in phosphate buffer for 1.5-2 hours depending on developmental stage, cryoprotected in sucrose, then embedded in Optimum Cutting Temperature (OCT) compound (Tissue-Tek, Torrance, CA) and stored at −80°C until sectioning. Tissue was cryosectioned sagittally at 12μm, and stored at −20°C until stained. Frozen sections were permeabilized by incubating in 0.1% Triton/PBS (PBT), and incubated in 3% normal donkey serum (NDS)/3% bovine serum albumin (BSA)/0.1% PBT for one hour at room temperature. The sections were then incubated with primary antibodies overnight at 4°C.

*Immunohistochemistry* was conducted using the following antibodies: rabbit anti-Sox2 (1:200, Santa Cruz), guinea pig anti-RBPMS (1:500, PhosphoSolutions), and rabbit anti-DCX (1:1000, Abcam). Species-specific AlexaFluor 488-, 568-, 647-conjugated secondary antibodies were used to detect primary antibodies (1:1500, Invitrogen). Following antibody staining, nuclei were counterstained with DAPI (4’,6-diamidino-2-phenylindole; 1:1000, Sigma). Following washes, slides were mounted (DakoCytomation fluorescent mounting medium) and imaged using a Zeiss AxioObserver fluorescence microscope equipped with an Apotome 2, 10x and 20x objectives, and Hamamatsu Orca Flash v4.0 digital camera. Images were processed using Zen software (Zeiss) and Photoshop CS6 (Adobe, San Jose, CA).

*EdU birth dating studies*. Dams were injected intraperitoneally with 30 mg/kg body weight of EdU (Invitrogen) at E14.5 and sacrificed at E18.5. Sections were immunostained using the Click-It Kit according to the manufacturers protocol (Invitrogen).

### Image sampling, quantification, and statistics

For analysis of immunostaining markers and EdU labeling, counting frames (GCL: 50μm x 150μm/50μm x 25μm, INL/IPL: 25μm x 150μm) were randomly placed in respective layers of interest using ImageJ (FIJI)^28^. At least three counting frames were analyzed per histological section, with at least three histological sections of the retina taken from 3-6 different animals for each stain. Cell counts were normalized to total DAPI+ cells, ensuring consistency between WT and cKO sections. Cell counts for immunostains and EdU were normalized as a percentage of total DAPI+ cells. The ‘integrated density’ plugin was used to quantify RBPMS staining in the GCL due to the cytosolic localization of the stain. To maintain consistency when using the integrated density measure, images were set to equal brightness and contrast in ImageJ and then converted to ‘binary, which captured either the presence or absence of a signal, disregarding any intensity differences in the signal. For analysis of DCX staining and morphometric analysis, cell width and length were analysed using a ‘X/Y’ ratio, respectively, similar to the minor and major axis measurements described by Yu et al. (2013)^24^. The ‘straight’ tool in ImageJ was used to measure retinal thickness and the distance travelled by migratory neuroblasts from the apical surface, or retinal pigment epithelium.

To ensure consistency among samples, cell counts were restricted to images taken from approximately the same region of the retina, as identified by retinal thickness. In all experiments, 3-5 Mllt11 cKO and WT embryos were used for quantifications using the unbiased and systematic sampling method described previously^17^. Bar charts and statistical testing were conducted using Graphpad Prism V9 software, with results shown as mean ± SD. In all quantification studies, statistical differences were determined with Welch’s t-test (two-tailed) with a significance level set at P≡0.05 (*P≤0.05, **P≤0.01, ***P≤0.001, ****P≤0.0001).

## ACKNOWLEDGMENTS

We gratefully acknowledge funding support from the Canadian Institutes of Health Research (CIHR PJT-388914). We thank Sarah Whitehead for assistance with animal husbandry. We also thank Dr. William Baldridge for sharing RBPMS antibodies and comments to the manuscript.

